# Soil nutrient adequacy for optimal cassava growth, implications on cyanogenic glucoside production: a case of konzo-affected Mtwara region, Tanzania

**DOI:** 10.1101/464594

**Authors:** Matema L.E. Imakumbili, Ernest Semu, Johnson M.R. Semoka, Adebayo Abass, Geoffrey Mkamilo

## Abstract

Soils in areas affected by konzo (a cassava cyanide intoxication paralytic disorder) are predominantly infertile and probably unable to supply cassava the nutrients it needs to achieve optimal growth. The soil nutrient levels in these areas, could also be influencing cyanogenic glucoside production in cultivated cassava, however there is hardly any knowledge on this. An assessment of soil nutrient levels on cassava fields in konzo-affected areas was therefore carried out to determine their adequacy for optimal cassava growth and how this influences cassava cyanogenic glucoside production. Konzo-affected Mtwara region, in Tanzania, was used as a case study area. Correlations between total hydrogen cyanide (HCN) levels in cassava roots and various soil nutrient levels on cassava fields were carried out and relationships between cyanide intoxication and soil nutrient levels on fields from which toxic cassava roots had been harvested were also investigated. The results showed that cassava grows under conditions of severe nutrient stress in the region. Soil nutrients found to be deficient on most fields, like potassium (mean = 0.09, SD = 0.05 cmol/kg), magnesium (mean = 0.26, SD = 0.14 cmol/kg) and zinc (mean = 1.34, SD = 0.26 mg/kg), are known to reduce cyanogenic glucoside levels in cassava roots when adequate in soils. Cyanogenic glucoside levels in cassava roots however increased by high levels soil phosphorous (r_s_ = 0.486, p = 0.026 for all varieties) and sulphur (r_s_ = 0.593, p = 0.032 and r_s_ = 0.714, p = 0.047; for bitter and sweet cassava varieties, respectively) on these soils. The likelihood of cassava cyanide intoxication was also increased on fields with high pH and iron levels. High levels of sulphur and phosphorus, to very high levels of iron occurred on some fields. How soil nutrient supply influences cassava cyanogenic glucoside production in the konzo-affected areas was established.

## Introduction

Infertile soils are unable to supply crops growing on them, with the nutrients they need to attain optimal growth. This greatly limits the yields of crops. Yield reductions are brought about by plant nutrient stress resulting from an inadequate supply of nutrients. Besides limiting yields, low soil fertility further affects the nutritional composition of crops, hence altering their nutritional quality [1]. Soils with low fertility, are able to reduce yields of hardy crops like cassava (*Manihot esculenta* Crantz) and soil fertility (low or high), can also influence cyanogenic glucoside production in cassava [2–4]. Cyanogenic glucosides are an important nutritional quality determining factor for cassava, as they determine its safe consumption. The ingestion of high levels of cyanogenic glucosides from fresh cassava roots, or from products produced from such roots, exposes humans to cyanide intoxication.

Due to cyanogenic glucosides, cassava has been many at times associated with cyanide intoxication in some rural communities where it constitutes the main component of the people’s diet. The cyanide intoxication cases often resulted in a health disorder called konzo (spastic paraparesis), which causes an irreversible paralysis of legs in affected individuals [5,6]. A number of Sub-Saharan African countries have been affected by konzo, and the disorder is reported as persistent in very deprived areas of Mozambique, the Democratic Republic of Congo (DRC), Tanzania [7], Central African Republic [8,9] and in eastern Cameroon [10,11]

A characteristic common to all areas affected by konzo is their low soil fertility. Except for konzo-affected areas in DRC and Cameroon, all other affected areas are situated along the coast or along a lake shore, they include; Nampula and Zambézia districts in Mozambique [12], Newala and Mtwara districts in Tanzania [13] and the lake shore Tarime district in Tanzania [14]. All mentioned areas predominantly have sandy soils, which have very low soil fertility [14–19]. Although not coastal, konzo affected areas in Bandundu Region in DRC are located in the Savannah zone, which primarily consists of relatively infertile sandy soils [19,20]. Soils in the non-coastal Eastern region of Cameroon are additionally described as having a sandy clay texture and as being largely degraded [21]. Soils in konzo-affected western Central African Republic, bordering the eastern region of Cameroon [10], are probably just as degraded and unable to be cropped. Being predominantly sandy, soils in konzo affected areas are also likely to be prone to water stress, because of the low water retention of sandy soils. Low soil fertility is however an additional characteristic of sandy soils.

The agronomic factors that lead to increased cyanogenic glucoside levels and exacerbate cyanide intoxication during konzo outbreaks include; farming systems dominated by highly toxic bitter cassava varieties and drought or dry season related water stress [12,13,22] (Fig 1). While the sandy nature of soils in most konzo-affected areas contributes to water stress and the associated increase in cyanogenic glucoside levels in cassava, the additional influence of nutrient supply, from these predominantly nutrient poor soils, cannot be ignored and needs to be investigated. A study was hence carried out to assess the adequacy of nutrient levels in soils for optimal cassava growth, in the konzo-affected areas and to determine how these soil nutrient levels affect cyanogenic glucoside production in cassava. Konzo-affected Mtwara region of Tanzania, was used as a case study.

**Fig 1.**
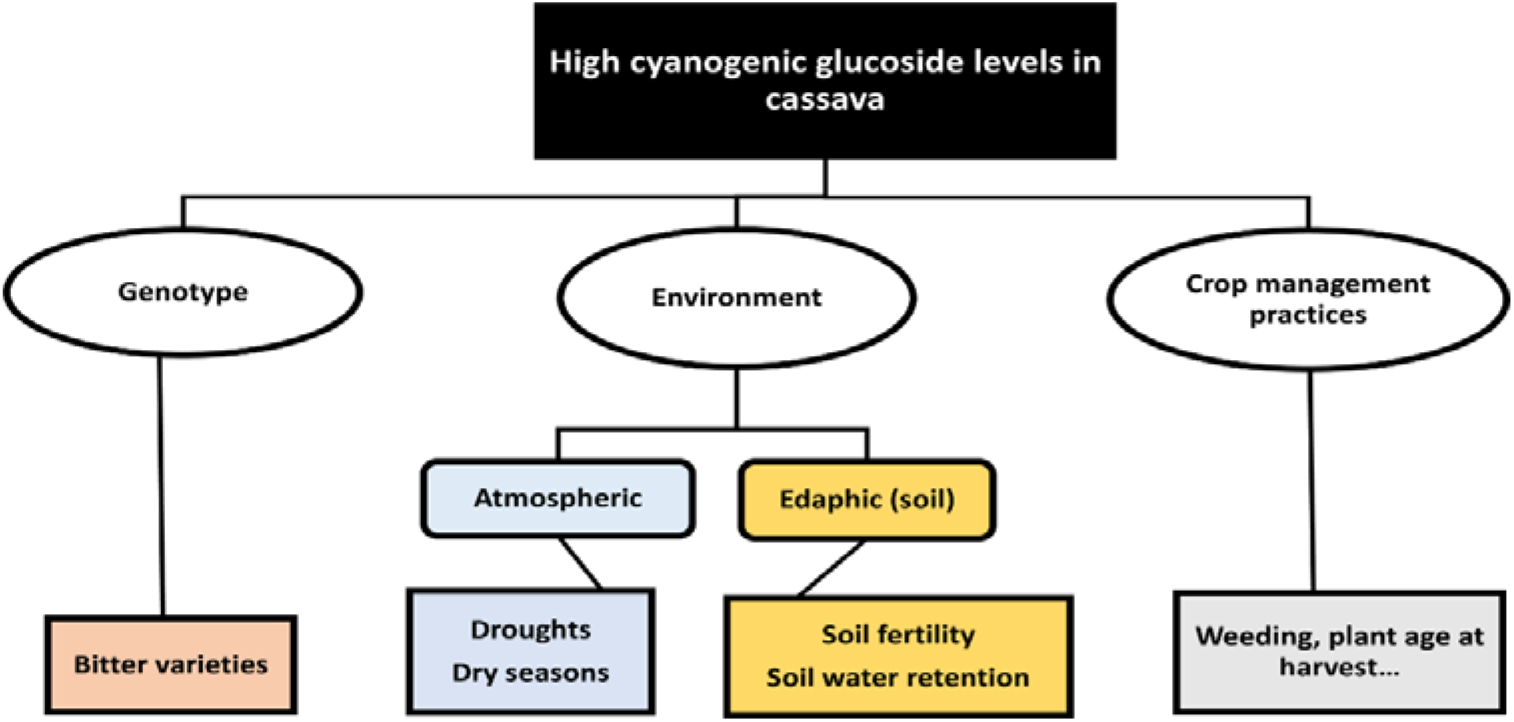
Agronomic reasons for high cyanogenic glucosides in cassava

## Materials and Methods

### Description of study area

The study was carried out in areas previously reported to have been affected by konzo in Mtwara region. Three districts have been reportedly affected by konzo in Mtwara region, namely Masasi, Mtwara Rural and Newala districts [13]. This study however focused on konzo-affected villages of Mtwara Rural and Newala districts, which are two of the five districts of Mtwara region (S 10°16′25″, E 40°10′58″). The two districts studied (Mtwara Rural and Newala) lie in Tanzania’s, Coastal Lowlands agroecological zone [17,23]. Soils in the region are generally classified as Ferralic Cambisols and are predominantly sandy [17,23]. The rainfall in the region is mono-modal and ranges from 800 to 1000 mm/year and the maximum and minimum temperatures vary from 29 to 31 °C and 19 to 23 °C, respectively [23]. Both districts lie on the Makonde plateau, with Newala lying to the west and Mtwara Rural to the eastern side of the plateau.

### Selection of households for the survey

The survey was carried out from 7^th^ to 16^th^ October, 2014, during the hot-dry season, which is a common harvest time for cassava in the region. The survey was carried out in some konzo affected villages of Newala district and Mtwara Rural district. The villages were selected from the 18 villages visited during a konzo rehabilitation and prevention program that was carried out in 2008 to 2009, by the Tanzania Food and Nutrition Centre (TFNC) and the Tanzania Red Cross Society (TRCS), with technical support from the Australian National University (ANU) [13]. Four villages were selected from each district, using simple random sampling. The villages from each district were numbered and randomised using the ‘RAND’ function in Microsoft Excel. The first 4 villages that appeared for each district in the randomised list were then picked. Mdimba, Ngalu, Songambele and Mkunjo were the villages selected from Newala district, whereas the villages Njengwa, Nyundo, Niyumba and Kiromba were selected from Mtwara Rural district.

Once again, using simple random sampling, 15 households were randomly picked from each village to participate in the survey. A 2012 village census list was used to select the households. The census list was the most recent of the past census carried out nationwide in Tanzania. Using the census list, each household in a village was given a unique number. The numbered list was then randomised using the ‘RANDBETWEEN’ function in Microsoft Excel and the first 15 households on the randomised list were picked. A total of 120 households were selected, 61 from konzo affected villages of Newala district and 59 from konzo-affected villages of Mtwara Rural district. Verbal consent was obtained from the household heads (farmers) before they could participate in the survey. If both spouses were present at a household, either spouse participated in the survey.

Prior to the survey, written permission to conduct the survey was sought from the government regional administrative offices of Mtwara region. The permission given was in letter form. This letter was then presented to the district offices in Newala and Mtwara rural districts. After obtaining verbal acknowledgement of the research from the district offices, the letter from the regional administrative office was then presented to the respective village administrative committees, who gave verbal consent for the work to proceed. A letter of introduction to present to the regional administrative office had been initially obtained from Sokoine University of Agriculture, which was the host institution for the researcher. This letter introduced the researcher by stating their name, the institution which they belonged to and the exact nature of the research to be carried-out. A humble request for the researcher to be supported was also included. The letter of introduction had been presented to the government regional administrative offices of Mtwara region.

### Data collected

In order to investigate soil nutrient adequacy and the implications that the soil nutrients have on cyanogenic glucoside production in Mtwara region, both soil and root samples were collected from cassava fields located in the konzo-affected villages of Mtwara Rural and Newala districts. Soil samples had however been only collected from 112 cassava fields and cassava roots from only 21 cassava fields. This is because there was a failure to collect soil and cassava root samples from all cassava fields belonging to the 120 households selected for the survey. Soil and root samples were collected and analysed as described in the sections below.

### Soil sample collection and analyses

Composite soil samples of about 500 g were collected from each household’s cassava field. This was done by collecting soil from at least 10 randomly selected points on the cassava fields from the top 20 cm layer of the soil [24]. Soil samples were first air-dried and later analysed for the following various soil chemical characteristics; soil pH, organic carbon (OC), total nitrogen (N), available phosphorous (P), exchangeable potassium (K), calcium (Ca) and magnesium (Mg), available sulphur (S), available zinc (Zn), copper (Cu), iron (Fe) and manganese (Mn) and exchangeable aluminium (Al). All laboratory methods used were as follows [25]: Soil pH was determined in a 1:1 soil to water solution and the Bray No. 1 method was used to extract P. The exchangeable cations K, Ca and Mg were extracted using 1 N ammonium acetate (NH_4_OAc) leaching solution (buffered at pH 7) and monocalcium phosphate [Ca(H_2_PO_4_)_2_] extracting solution was used to extract S. Zinc, Cu, Fe and Mn were extracted using diethylenetriaminepentaacetic acid (DTPA), N was determined using the micro Kjeldhal method and OC using the Walkley and Black dichromate method. Exchangeable Al was extracted using 1 N potassium chloride (KCl) extracting solution and it was only carried out on soil samples that had a pH of less than five. The value obtained for exchangeable Al and the respective Ca, Mg and K levels for the soils were used to calculate the soils’ Al saturation, using Equation 1 [24].

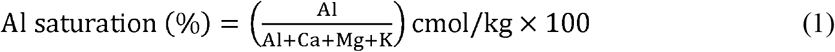

The soil texture of the sampled soils had also been determined. This has been done using the hydrometer method.

### Root sampling and total hydrogen cyanide determination

To assess root cyanogenic glucoside content, root samples of commonly grown bitter and sweet cassava varieties in the konzo-affected villages were collected from the cassava fields where the soil samples had been collected. Cassava root samples were collected from four randomly selected plants of the same variety from a farmers’ field. Only plants identified by famers as being ready for harvest were selected. Three roots were sampled per plant for hydrogen cyanide (HCN) analysis (that is, potential cyanogenic glucoside determination) [26]. The total HCN content of fresh cassava roots was determined within 24 hours after harvest using the picrate paper method [27,28]. A 100 mg section of fresh cassava root taken from the middle of the root was placed in a vial with buffer solution and a picrate paper. The contents in the vial were then left to incubate in the dark at room temperature for 16 to 24 hours. The picrate papers darkened by the cyanide liberated from the fresh root sections, were then eluted in 5 ml of distilled water. The absorbance of the solution so obtained was then determined using a spectrophotometer at 510 nm. Cassava root HCN content on a fresh weight basis was then calculated using Equation 2.

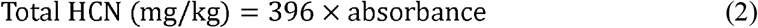

where 396 is a value derived from the calibration curve between picrate solutions placed in known standard cyanide solutions and their absorbance values [27,28].

### Farmer interviews

To further investigate the influence of soil nutrient supply on cyanogenic glucoside production in konzo-affected areas, links between soil nutrient supply and cases of cassava cyanide intoxication from cassava roots harvested on these soils were also investigated. Individual farmer interviews were carried out to obtain information on households that had experienced cassava cyanide intoxication. Farmers were asked whether any of their household members had experienced; dizziness, headaches, stomach pains, nausea, vomiting, brief confusion, difficulty in standing, difficulty in speaking (a heavy tongue) or even death after consuming cassava [29,30]. Clinical validation of symptoms experienced had not been obtained, the responses were hence based on the farmers understanding of cassava cyanide intoxication. All farmers appeared to be aware of cassava cyanide intoxication and its associated symptoms.

Farmers who agreed to have had at least one household member affected by cyanide intoxication, after consuming cassava, were also asked whether the cassava roots that had brought about the intoxication had been obtained from the households’ own cassava field. For the cyanide intoxication cases to be linked to soil nutrient levels on fields from which the poisonous cassava roots had been harvested, farmers were also asked whether they still cultivated the field(s) from which the toxic cassava roots had been harvested. Soil samples were then collected from the respective field or fields cropped at the time of the intoxication experience. Soils from cassava fields of households not affected by cassava cyanide intoxication were also collected and analysed as described in a previous section.

### Data Analysis

Soil nutrient levels and other important soil chemical characteristics on cassava fields were analysed using descriptive statistics (means, standard deviations, and minimum and maximum values). The adequacy of soil nutrient levels for cassava production was assessed using critical levels and sufficiency ranges of nutrients known to be suitable for cassava production. Root HCN levels of collected root samples were also analysed using descriptive statistics. As the data was not normally distributed, the Spearman’s correlation (two-tailed) was used to assess the statistical significance of relationships between cassava root HCN levels and various soil nutrient levels. The influence that soil nutrient supply has on causing cassava root toxicity and associated cases of cassava cyanide intoxication was also examined using the binomial logistic regression analysis. The hypothesis tested was that “the likelihood of cassava cyanide intoxication occurring was related to soil nutrient levels on crop fields on which the toxic roots were harvested.” All statistical procedures were carried out as outlined by [31] and all analyses were carried out using IBM SPSS Statistics, version 20.

## Results and Discussion

### Suitability of soils for optimal cassava growth

Table 1 shows the soil nutrient levels on cassava fields in the konzo-affected areas together with their suitability for optimal cassava growth. Levels of other important soil chemical characteristics like pH, OC and Al saturation, are also included in Table 1.

**Table 1.** Soil chemical characteristics of soils on cassava fields in konzo-affected villages of Mtwara Region and their suitability for cassava production

| Soil parameter | Mean<br>n = 112 | Standard |  |  |  |  | Range/level rated as suitable for cassava<br>production | Rated according to: |
| --- | --- | --- | --- | --- | --- | --- | --- | --- |
|  |  | deviation | Minimum |  | Maximum |  |  |  |
|  |  | SD | Value | Rating | Value | Rating |  |  |
| pH | 5.29 | 0.56 | 4.40 | l | 6.96 | m | 4.5 – 7.0 | [24] |
| OC (%) | 0.63 | 0.27 | 0.31 | vl | 1.22 | vl | 4.0 – 10.0 | [32] |
| N (%) | 0.07 | 0.03 | 0.01 | vl | 0.33 | m | 0.20 – 0.50 | [32] |
| P (mg/kg) | 8.21 | 7.62 | 1.19 | l | 34.52 | h | < 4.2 | [33] |
| K (cmol/kg) | 0.09 | 0.05 | 0.02 | vl | 0.32 | h | 0.15 – 0.25 | [24] |
| Ca (cmol/kg) | 2.02 | 0.93 | 0.46 | l | 4.49 | m | 1.0 – 5.0 | [24] |
| Mg (cmol/kg) | 0.26 | 0.14 | 0.02 | vl | 0.76 | m | 0.40 – 1.00 | [24] |
| S (mg/kg) | 5.30 | 3.17 | 0.60 | l | 23.51 | h | < 6.0 | [32] |
| Zn (mg/kg) | 0.34 | 0.26 | trace | vl | 1.79 | m | 1.0 – 3.0 | [34] |
| Cu (mg/kg) | 0.01 | 0.05 | trace | vl | 0.43 | m | 0.3 – 0.8 | [34] |
| Fe (mg/kg) | 32.87 | 19.10 | 3.19 | l | 83.45 | vh | 4.0 – 6.0 | [34] |
| Mn (mg/kg) | 9.74 | 10.03 | 1.29 | m | 71.71 | vh | 1.2 – 3.5 | [34] |
| Al Saturation (%)□ | 9.27 | 9.19 | 0.00 | m | 32.94 | m | < 75.0 | [24] |

Where: vl, l, m, h and vh stand for very low, low, medium, high and very high.
□ n = 72 for Al saturation.
The soil texture classes for the sampled soils are as follows; 58.0% Loamy sand, 22.3% Sandy clay loam, 13.4% Sandy loam and 6.3% Sand.

Soil pH levels in the region were mainly in the desirable range and were hence not too low or too high for good cassava growth (Tables 1). Only 0.9% of the fields had soil pH levels below 4.5, while none had pH levels above 7.0. The pH of soils in Mtwara region is not as low as might be expected for highly weathered soils and is usually > 5.0 because of the presence of limestone in the parent material of these soils [35]. Limestone has reducing effects on soil pH. Aluminium toxicity similarly posed no limitations on cassava growth with all fields having none or low levels of Al saturation.

Almost all (99.1%) the fields sampled had N levels below what is considered as sufficient for a broad range of tropical crops [32], including cassava (Table 1). Soil organic carbon is a main source of N for crops grown without fertilizer application. Fertiliser use is low amongst these rural poor farmers, the low OC levels, which were all less than 2% in all the fields’, hence explains the low levels of N in these soils. In regard to cyanogenic glucoside production, an improved supply of N, on N deficient soils, was able to reduce root HCN levels in one study [36]. Improving the supply of N could thus be beneficial for reducing cassava root HCN levels on these N deficient soils. High levels of N are however mainly associated with increasing cyanogenic glucoside levels in cassava plants [3].

Only 34.8% of the sampled fields had soil P levels below the critical level of 4.2 mg/kg. Soil P levels varied from very low to high on the sampled fields, with more fields having sufficient P (Table 1). Not much research has been done on the relationship between soil P availability and root HCN accumulation. One study however concluded that P had no influence on cassava root HCN levels [37]. When P is adequate in soils, root development is greater, enabling greater water uptake particularly during less moist periods. With adequate P, cassava plants would thus be able to mitigate water stress and the associated increase in cyanogenic glucoside levels in cassava roots.

Another important nutrient limitation in the sampled soils was low soil K. About 84.3% of the sampled fields had soil K levels below the sufficiency range considered adequate for healthy cassava growth. An adequate supply of soil K, through fertilizer application has been shown to often reduce HCN levels in cassava roots [3,38,39]. Some studies have however shown no effects on root HCN with an improved supply of K [40]. The low soil K levels on these cassava fields could thus contribute to increased root cyanogenic glucoside levels in cassava produced in the region.

The nutrients Ca, Mg and Zn were deficient in 13.9%, 84.3% and 93.0% of the fields, respectively. In most cassava fields, levels of soil Ca were mainly adequate for cassava production. Like pH, adequate Ca can be attributed to the presence of limestone in the parent material of these soils. Reduced root HCN levels have been reported with improved Ca supply in soils [36,41]. Unlike Ca, Mg and Zn were more severely deficient in most fields. An increased supply of the nutrients Mg and Zn in soils, has been shown to reduce cassava root HCN levels [3,36]. Furthermore, reported reductions in cassava root HCN levels, with the application of ash, are attributed to the presence of K, Ca and Mg in the ash. Deficiencies of K, Ca, Mg and Zn could thus result in increased cyanogenic glucoside production in cassava produced in konzo-affected areas in Mtwara region.

Sulphur was deficient in 63.5% of the sampled fields. Reductions in root HCN levels with an increased supply of soil S have been reported in some studies [36,41] implying that low levels of S could have an increasing effect on root cyanogenic glucoside production in Mtwara region. Unlike S, Cu was deficient on almost all (95.7%) of the sampled fields, whereas Fe and Mn were mainly sufficient to very high in all fields (Table 1). High levels of Fe and Mn are expected as soils in Mtwara region are predominantly Ferralic cambisols [17,23] and are thus dominated by Fe and Mn oxides and hydroxides [42]. None of the fields had low Mn levels and only 0.9% of the fields had low soil Fe levels. There are hardly any reports on the influence of Cu, Fe or Mn on cassava root HCN levels. The very low soil Cu levels could however cause nutrient stress in cassava grown in the region. There is also a possibility for the very high levels of Fe and Mn in these soils to negatively influence cyanogenic glucoside production in cassava produced in the konzo-affected areas of Mtwara region. Very high soil nutrient levels are toxic to cassava plants and can cause yield reductions [43], they can also probably result in increased cyanogenic glucoside levels.

### Relationships between soil nutrients and root cyanogenic glucoside content

The results of the Spearman’s correlations, carried out to determine relationships between soil nutrient levels and root HCN levels of cassava, in the konzo-affected villages, are shown in Table 2. Correlations between soil nutrients and root HCN levels of bitter and sweet cassava varieties were also separately assessed.

**Table 2.** Correlations between root HCN levels and various soil properties

| Soil parameter | Bitter varieties<br>(n = 13) |  | Sweet varieties<br>(n = 8) |  | All varieties<br>(n = 21) |  |
| --- | --- | --- | --- | --- | --- | --- |
| | $r_s$ | p - value | $r_s$ | p - value | $r_s$ | p - value |
| pH | 0.135 | 0.660 | -0.407 | 0.317 | 0.072 | 0.758 |
| OM | 0.482 | 0.095 | 0.675 | 0.066 | 0.149 | 0.519 |
| N | -0.208 | 0.496 | 0.182 | 0.666 | -0.023 | 0.922 |
| P | 0.500 | 0.082 | 0.238 | 0.570 | **0.486** | 0.026 |
| K | -0.394 | 0.183 | -0.443 | 0.272 | -0.106 | 0.649 |
| S | **0.595** | 0.032 | **0.714** | 0.047 | 0.374 | 0.095 |
| Ca | 0.014 | 0.964 | -0.072 | 0.865 | 0.100 | 0.666 |
| Mg | -0.339 | 0.257 | 0.132 | 0.756 | 0.018 | 0.940 |
| Zn | 0.262 | 0.387 | 0.491 | 0.217 | 0.356 | 0.113 |
| Cu | . | . | -0.082 | 0.846 | -0.222 | 0.334 |
| Fe | -0.157 | 0.609 | 0.333 | 0.420 | -0.014 | 0.951 |
| Mn | -0.225 | 0.459 | -0.476 | 0.233 | -0.001 | 0.996 |
| Al Saturation | -0.152 | 0.774 | -0.100 | 0.873 | -0.116 | 0.733 |

Correlation coefficients ( $r_s$ ) in bold are significant at $p < 0.050$ using the Spearman's correlations (two-tailed)

Statistically significant positive correlations were found between soil S and root HCN levels of both sweet and bitter cassava varieties (Table 2). The positive correlation coefficient, indicated that root HCN levels increased with soil S in both bitter and sweet cassava varieties. Another statistically significant positive correlation was found, this time between root HCN levels of all sampled cassava varieties (bitter and sweet combined) and soil P levels on cassava fields. This showed that root HCN levels in all varieties generally increased with increased P in these soils.

The results are contrary to the previously mentioned findings by other studies, on the effects of improved soil S and P levels on cassava root HCN levels. It is however important to note that some of the soils in the region had high levels S and P that were outside the range suitable for cassava optimal cassava growth (Table 1). Although not toxic (very high), these soil S and P levels may still cause nutrient stress. The results indicate that cassava grown in the konzo-affected villages could be more sensitive to high S and P levels than to low levels of these nutrients. A study, carried out under controlled conditions, had however failed to establish relationships between soil N and HCN levels in cassava roots and leaves [44]. It is thus possible that cassava cyanogenic glucoside content is not always well correlated to soil nutrient supply, hence the observed results.

### Cyanogenic glucoside content of sampled cassava varieties

Root HCN levels for the varieties sampled from the konzo-affected villages in Mtwara region are given in Table 3.

**Table 3.** Total hydrogen cyanide levels of roots of various cassava varieties from konzo-affected Mtwara region

| Variety | Type | n | HCN levels, fresh weight basis (mg/kg) |  |  |
| --- | --- | --- | --- | --- | --- |
|  |  |  | Minimum | Maximum | Mean |
| *Badi* | Sweet | 1 | 16.8 | . | . |
| *Kigoma* | Sweet | 4 | 43.1 | 58.1 | 49.5 |
| *Mnalile Kuchumba* | Sweet | 3 | 25.9 | 109.8 | 62.2 |
| *Limbanga* | Bitter | 2 | 114.4 | 181.9 | 148.2 |
| *Mohammed Mfaume* | Bitter | 1 | 79.4 | . | . |
| *Musa Saidi* | Bitter | 3 | 51.0 | 131.1 | 85.15 |
| *Namanjele* | Bitter | 1 | 44.3 | . | . |
| *Nanjenjeha* | Bitter | 3 | 54.3 | 144.3 | 98.1 |
| *Salanga* | Bitter | 3 | 111.3 | 191.9 | 147.5 |

All root samples of the bitter cassava varieties, *Limbanga* and *Salanga*, had HCN levels above 100 mg/kg (Table 3); these levels exceeded 50 mg/kg, which is the limit set for the safe consumption of fresh cassava roots [45]. The two varieties thus had a great potential of causing cyanide poisoning. Some root samples of the bitter varieties *Nanjejeha* and *Namanjele* were however below 50 mg/kg. This showed that some bitter cassava varieties can also at times contain lower HCN levels and may not always be toxic. Root samples of the sweet cassava varieties, *Mnalile Kuchumba* and *Kigoma*, had total HCN levels above 50 mg/kg. This showed that sweet cassava varieties, produced in the konzo-affected areas, are also capable of accumulating high levels of cyanogenic glucosides. It is important to note that all root samples had been harvested during the same period; differences in root HCN levels due to differences in atmospheric conditions (seasonal influence) were hence eliminated.

### Relationships between soil nutrient levels on cassava fields and cyanide intoxication from cassava harvested from them

Out of the 120 farmers interviewed 45.8% mentioned that they had at least one household member affected by cassava cyanide intoxication. The intoxication experiences were either acute or they could have resulted in konzo or even death. Some farmers could remember the exact year in which the cyanide intoxication occurred, this information in shown in Table 4. While some cases occurred more than 30 years ago, some cases of cassava cyanide intoxication were relatively recent.

**Table 4.**
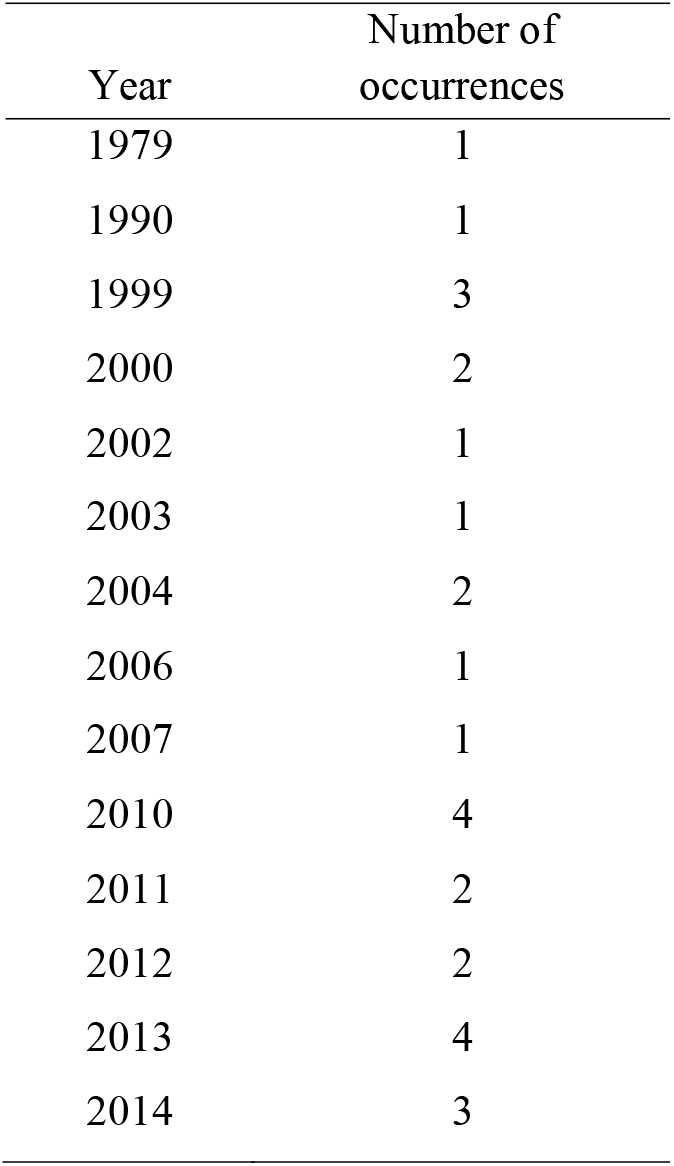
Cassava cyanide intoxication occurrences in konzo-affected areas of Mtwara region

| Year | Number of occurrences |
| --- | --- |
| 1979 | 1 |
| 1990 | 1 |
| 1999 | 3 |
| 2000 | 2 |
| 2002 | 1 |
| 2003 | 1 |
| 2004 | 2 |
| 2006 | 1 |
| 2007 | 1 |
| 2010 | 4 |
| 2011 | 2 |
| 2012 | 2 |
| 2013 | 4 |
| 2014 | 3 |

Only 23.2% of the farmers interviewed were still using the field, from which the cassava roots that caused intoxication had been harvested. The likelihood of a household getting a case of cyanide intoxication or not, because of their fields’ soil nutrient levels were investigated using logistic regression analysis. The results of the logistic regression analysis are shown in Table 5.

**Table 5.** Results of the logistic regression analysis showing the connection between cassava cyanide intoxication and soil nutrient supply in konzo-affected areas of Mtwara region

| Predictor | $\beta$ | SE | $X^2$ | df | p-value | Exp($\beta$) |
| --- | --- | --- | --- | --- | --- | --- |
| pH | 2.640 | 0.926 | 8.124 | 1 | 0.004 | 14.014 |
| OC | -1.754 | 2.504 | 0.491 | 1 | 0.483 | 0.173 |
| N | -0.980 | 14.205 | 0.005 | 1 | 0.945 | 0.375 |
| P | -0.071 | 0.056 | 1.603 | 1 | 0.205 | 0.931 |
| K | 1.867 | 7.139 | 0.068 | 1 | 0.794 | 6.468 |
| S | 0.090 | 0.092 | 0.973 | 1 | 0.324 | 1.095 |
| Ca | -0.061 | 0.539 | 0.013 | 1 | 0.910 | 0.941 |
| Mg | -0.446 | 3.425 | 0.017 | 1 | 0.896 | 0.640 |
| Fe | 0.062 | 0.026 | 5.740 | 1 | 0.017 | 1.064 |
| Cu | 11.745 | 6.898 | 2.899 | 1 | 0.089 | 126114.5 |
| Zn | 0.017 | 1.468 | 0.000 | 1 | 0.991 | 1.017 |
| Mn | 0.061 | 0.035 | 3.089 | 1 | 0.079 | 1.063 |
| Constant | -16.919 | 5.594 | 9.149 | 1 | 0.002 | 0 |
| Test | | | $X^2$ | df | p-value | |
| Overall model evaluation |  |  |  |  |  |  |
| Score test |  |  | 25.958 | 12 | 0.011 |  |
| Wald test |  |  | 27.937 | 1 | 0.000 |  |
| Goodness-of-fit test |  |  |  |  |  |  |
| Hosmer & Lemeshow |  |  | 7.588 | 8 | 0.475 |  |

Nagelkerke $R^2 = 0.316$ ; c-statistic = 81.1%. Statistical significance was determined at $p < 0.050$ .
Note: Al saturation was not included as a predictor variable, as only a few samples had a value for it.

The equation derived from the results of the logistic regression analysis are shown in Equation 3:

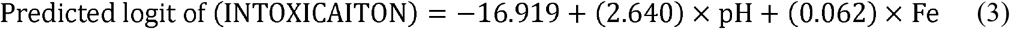

The model produced, showed that high pH contributed to the likelihood of cassava cyanide intoxication in these communities and so did high levels of Fe in soils. Cassava growing on fields with high soil pH levels and high Fe levels are hence more likely to produce toxic cassava roots. Although some fields had soil pH levels above 6, soil pH levels on these cassava fields did not exceed the upper limit (pH = 7) of the pH range recommended as suitable for optimal cassava growth (Table 1). The cassava varieties grown in these areas could hence be sensitive to soil pH levels within the optimal pH range recommended for cassava. The cassava varieties also appear to be better adapted to lower pH levels, in terms of low cyanogenic glucoside production. In agreement with these findings, cassava root bitterness, which is mostly correlated to root HCN levels, was reported to increase with soil pH (r = 0.339, p = 0.047) in soils with pH values ranging from 5.8 to 7.8 (1:2.5 soil to water) [46].

In the case of Fe, some cassava fields had very high soil Fe levels, making the observed results somewhat expected due to the toxic levels of Fe in these soils (Table 1). However, in contrast to the findings of the present study, significantly lower root HCN levels were reported in cassava grown on iron rich soils compared to levels found in cassava grown on fields with lower soil Fe levels [4]. The findings of the present study however agree with the farmers own perceptions of the causes of increased cassava root bitterness and its associated toxicity in these konzo-affected areas. The farmers perceived that root bitterness increased when cassava was grown on higher iron-containing red soils, in Mtwara region [47].

### Conclusion

Soils on cassava fields in konzo-affected areas of Mtwara Region are severely inadequate in multiple soil nutrients. The soils are thus unable to supply most nutrients in amounts needed to support optimal cassava growth. Increased cyanogenic glucoside production in cassava produced is expected, particularly from deficiencies of K, Mg and Zn, whose sufficiency in soils is known to reduce cyanogenic glucoside levels in cassava. Despite evidence of severely low soil fertility in the region, high cassava cyanogenic glucoside levels are however linked to high levels of P and S on these soils. Furthermore, increased cassava cyanide intoxication also occurs when cassava is cultivated on soils with high soil pH and Fe. High levels of S and P (which occurred on some cassava fields) in these soils, together with the high to very high levels of Fe (which was prevalent on most fields) could thus be contributors of high cyanogenic glucoside levels in konzo-affected areas of Mtwara region in Tanzania. Similar trends could also be occurring in other konzo-affected areas.

## Acknowledgements

The authors thank the farmers for willingly participating in the survey. They also thank the government administrative staff in Mtwara region for their role in ensuring that the study took place. The authors also thank the staff at Naliendele Agricultural Research Institute in Mtwara, for their assistance during the execution of the survey. Lastly, the authors thank the staff at the soil science laboratory at Sokoine University of Agriculture for their guidance and support during sample analysis.

